# Discovery of powdery mildew resistance gene candidates from *Aegilops biuncialis* chromosome 2M^b^ based on transcriptome sequencing

**DOI:** 10.1101/698324

**Authors:** Huanhuan Li, Zhenjie Dong, Chao Ma, Xiubin Tian, Zhiguo Xiang, Qing Xia, Pengtao Ma, Wenxuan Liu

## Abstract

Powdery mildew is one of the most widespread diseases of wheat. Breeding resistant varieties by utilization of resistance genes is considered as the most economic and effective method of controlling this disease. Previous study showed that the gene(s) at 2M^b^ in Chinese Spring (CS)-*Aegilops biuncialis* 2M^b^ disomic addition line TA7733 conferred high resistance to powdery mildew. In this study, 15 *Bgt* isolates prevalent in different regions of China were used to further test the resistance spectrum of TA7733. As a result, TA7733 was high resistance to all tested isolates, indicating that the gene(s) on chromosome 2M^b^ was broad-spectrum powdery mildew resistance. In order to mine resistance gene candidates and develop 2M^b^-specific molecular markers to assist the transfer resistance gene(s) at chromosome 2M^b^, RNA-seq of TA7733 and CS was conducted before and after *Bgt*-infection, generating a total of 158,953 unigenes. Of which, 7,278 unigenes were TA7733-specific which were not expressed in CS, and 295 out of these 7,278 unigenes were annotated as R genes. Based on Blastn against with CS Ref Seq v1.0, 61 R genes were further mapped to homoeologous group 2. Analysis of R gene-specific molecular markers designed from R gene sequences verified 40 out of 61 R genes to be 2M^b^ specific. Annotation of these 40 R genes showed most genes encoded nucleotide binding leucine rich repeat (NLR) protein, being most likely resistance gene candidates. The broad-spectrum powdery mildew resistance gene(s), disease resistance gene candidates, and functional molecular markers of 2M^b^-specific in present study will not only lay foundations for transferring disease resistance gene(s) from 2M^b^ to common wheat by inducing CS-*Ae. biuncialis* homoeologous recombination, but also provide useful candidates for isolating and cloning resistance gene(s) and dissecting molecular and genetic mechanisms of disease resistance from 2M^b^.

## Introduction

Common wheat (*Triticum aestivum* L., 2*n* = 6*x* = 42, AABBDD), one of the most widely planted crops in the world provides 20% of the calories and 25% of its protein consumed by human [1,2]. Wheat production plays an important role in food security and social stabilization. However, wheat yields and quality are seriously threatened by various diseases, such as rusts, *Fusarium* head blight (FHB) and powdery mildew. Wheat powdery mildew, caused by *Blumeria graminis* f. sp. *tritici* (*Bgt*), is one of the most destructive diseases all over the world, with severe yield losses ranging from 13% to 50% [3,4]. In recent years, certain agronomic practices to increase yields, such as popularization of high planting density, high inputs of irrigation and fertilization have accelerated the spread and severity of powdery mildew [5,6]. Though spraying fungicides can reduce the damage caused by this disease to some extent, it can also result in side effects such as drug resistant of powdery mildew fungus, environment pollution and high production cost [7]. Breeding disease resistant varieties is currently considered as the most effective and economical method to control powdery mildew.

Wild relatives of common wheat contain a large number of *Bgt* resistance genes for wheat improvement. Up to now, the number of designated genes of powdery mildew resistance was more than 80 at 54 loci [8,9], of which approximately half of the powdery mildew resistance genes (*Pm*) were derived from wild relatives of common wheat. However, some *Pm* genes had been defeated by new virulent races or by races that were previously present at very low frequencies in the pathogen population [10,11], and some were difficult to use in wheat breeding because of linkage drags [12,13]. So, ongoing efforts to explore and identify new powdery mildew resistance genes for wheat breeding programs are also needed.

*Aegilops biuncialis* (2*n* = 4*x* = 28, U^b^U^b^M^b^M^b^) is a wild relative of wheat, belonging to the section *Polyeides* of the genus *Aegilops*. The U^b^ genome of it was derived from *Ae. umbellulata* (2*n* = 2*x* = 14, UU), and M^b^ genome from the diploid *Ae. comosa* (2*n* = 2*x* = 14, MM) [14,15]. *Aegilops biuncialis* owns many important agronomic traits of wheat improvement need, such as resistance to yellow rust [16], brown rust [17], powdery mildew and barley yellow dwarf virus [18], tolerance to drought and salt [19-21], high micronutrients contents [22], and special high molecular weight glutenin subunits [23]. Successful attempts have been made to cross *Ae. biuncialis* with wheat, develop a series of wheat-*Ae. biuncialis* addition lines, and transfer desirable genes from *Ae. biuncialis* into wheat [20,24,25]. It was observed that CS-*Ae. biuncialis* 2M^b^ disomic addition line TA7733 conferred high resistance to powdery mildew compared with its recipient parent CS in our laboratory [26].

The isolation and cloning of plant disease resistance genes had great significance for both plant disease resistance breeding and the study on molecular mechanisms of disease resistance. Map-based cloning is currently an important method to isolate novel genes. Whereas, it is very challenging to perform fine mapping and map-based cloning of genes derived from wild relatives of wheat [27], because alien-wheat homoeologous recombinations are strictly under the control of *Ph* genes in hexaploid wheat backgrounds [28,29]. Furthermore, molecular markers of alien chromosome-specific were limited for fine mapping of alien genes. Regardless, with the rapid development of high-throughput sequencing, sequencing-based technologies such as RNA-seq have been frequently used to develop molecular markers [1,30,31], detect gene expression pattern and level responded to pathogens [32], exploit new genes and identify gene function without prior information of the particular reference genome [33,34]. RNA-seq is very helpful to explore disease-resistant genes (R) derived from wild relatives. For example, Li et al. (2016) obtained eight powdery mildew resistance related genes from *Thinopyrum intermedium* by RNA-seq analysis [35]. Zou et al. (2018) successfully isolated a powdery mildew resistance gene *Pm60* from *T. urartu* by combining genetic mapping and RNA-seq analysis [9].

In this study, we report the assays of a broad-spectrum resistance gene(s) on chromosome 2M^b^ derived from *Ae. biuncialis*, discovery of candidate genes of powdery mildew resistance, and development of molecular markers of chromosome 2M^b^ specific based on transcriptome sequencing of CS-*Ae. biuncialis* 2M^b^ disomic addition line TA7733 applied before and 12-72 h post *Bgt* isolates inoculation. These will provide the foundations for transferring and cloning resistance gene(s) from chromosome 2M^b^ as well as further understanding the molecular and genetic mechanisms of disease resistance.

## Materials and Methods

### Plant materials

Common wheat landrace CS (2*n* = 6*x* = 42, AABBDD), *Ae. comosa* TA2102 (2*n* = 2*x* = 14, MM), and CS-*Ae. biuncialis* 2M^b^ disomic addition line TA7733 (2*n* = 44) were used in this study. All the materials were kindly provided by the Wheat Genetics Resource Center at Kansas State University, and maintained at the experimental station of Henan Agricultural University, China.

### Cytogenetic analysis

Chromosome spreads were prepared from root tip cells as described by Huang et al. (2018) [36]. The cytological observations were performed using a BX51 Olympus phase contrast microscope (Olympus Corporation, Tokyo, Japan).

Genomic DNA was extracted from fresh leaves using a modified CTAB (hexadecyl trimethyl ammonium bromide) method [37]. The concentration and purity of DNA were measured with the Nanophotometer P360 (Implen GmbH, München, Germany).

GISH was applied to analyze the chromosomal composition of TA7733. Genomic DNA of *Ae. comosa* accession TA2102 (genome M^b^ donor of *Ae. biuncialis*) and wheat CS were respectively used for probe labeling with fluorescein-12-dUTP and blocking at a ratio of 1:130 to distinguish *Ae. biuncialis* 2M^b^ chromosome. GISH was carried out as described by Liu et al. (2017) [38]. Hybridization signals were observed under an OLYMPUS AX80 (Olympus Corporation, Tokyo, Japan) fluorescence microscope, captured with a CCD camera (Diagnostic Instruments, Inc., Sterling Heights, MI, USA) and processed with Photoshop CS 3.0.

After GISH, the hybridization signals were washed off with PBS (phosphate-buffered saline). Eight single-strand oligonucleotides were then used as probes for dual-color ND-FISH [36,39]. The eight oligonucleotides includes Oligo-pAs1-1, Oligo-pAs1-3, Oligo-pAs1-4, Oligo-pAs1-6, Oligo-AFA-3, Oligo-AFA-4, Oligo-pSc119.2-1 and Oligo-(GAA)_10._ The first six, were labeled with 6-carboxytetramethylrhodamine (TAMRA) generating red signals, and the last two being labeled with 6-carboxyfuorescein (FAM) generating green signals. All the oligonucleotides were synthesized at Sangon Biological Technology, Shanghai, China.

### Evaluation of powdery mildew resistance

A mixture of prevailing *Bgt* isolates collected in Henan Province were used to evaluate the resistance of CS and CS-*Ae. biuncialis* 2M^b^ disomic addition line TA7733. Fifteen *Bgt* isolates prevalent in different regions of China were chosen to evaluate the resistance spectrum of TA7733 at the seedling stage. The infection type (IT) were scored 7-10 days post inoculation using a 0 to 4 rating scale [40], with 0 as immune, 0; as nearly immune, 1 as highly resistant, 2 as moderately resistant, 3 as moderately susceptible, and 4 as highly susceptible. IT 0 to 2 were considered as resistance, while IT 3 to 4 were being susceptible.

At 10 days post *Bgt* inoculation, the first leaves of TA7733 and CS were cut into 2 cm segments and stained with coomassie brilliant blue following Li et al. (2016) [35].

### Illumina library construction and sequencing

Seeds of CS and TA7733 soaking in water for 24 h at 23°C were transferred into a mixture of nutrient soil and vermiculite (1:1). Seedlings with full extended first leaf were dusted using fresh conidiophores of *Bgt* isolates. Leaves at 0, 12, 24, 48 and 72 hours post-inoculation (hpi) were respectively collected, rapidly frozen in liquid nitrogen and stored at −80°C for RNA extraction.

Total RNA of ten samples (five time points post inoculation for CS and TA7733, each) were extracted for transcriptome sequencing. Then mixture of equal amounts of RNA, at 12, 24, 48 and 72 hpi from TA7733 and CS were used as RNA-seq sample RI and SI, respectively. RNA at 0 hpi from TA7733 and CS were accordingly represented as RC and SC. Four libraries with an insert size of 200 bp were constructed from these RNA samples (RI, RC, SI and SC), and then sequenced using the Illumina HiSeqTM 2500 by the Beijing Genomics Institute.

### Reads processing, assembly and sequence annotation

Prior to assemble, sequencing raw reads were pre-processing using a Perl script dynamic-Trim.pl to remove the adaptor sequences, low-quality sequences, low complexity sequences, short reads and empty reads. Reads data with a quality score (Qphred) ≥ 50 (Q50: ratio of an error rate 0.01%) were then merged and input into the data assembly software Trinity for assembling into transcripts. The generated unigenes were annotated by a Blastx alignment search (E-value<10^-5^) against the NCBI non-redundant (NR) protein, SWISSPROT, gene ontology (GO), eukaryotic orthologous groups (KOG), kyoto encyclopedia of genes and genomes (KEGG) and plant resistance gene (PRG) databases.

### Amplification and analyses of candidate disease resistance genes

Sixty-one pairs of unigene-specific primers were used to identify the different unigenes using PCR. PCR amplification were conducted in 15 μl reactions containing 2 μl template gDNA (100 ng/µl), 0.25 µl forward primer (10 µmol/l), 0.25 µl reverse primer (10 µmol/l), 7.5 µl Taq MasterMix (CW Bio Inc., China) and 5 µl ddH_2_O. PCR cycling conditions were as follows: 94°C for 5 min followed by 35 cycles of 94°C for 30 s, 52-66°C for 30 s, and 72°C for 1 min, followed by a final 10-min extension at 72°C. The amplified PCR products were separated on a 2.0% agarose gel-electrophoresis stained with ethidium bromide and visualized by UV light.

### Mapping candidate disease resistance genes onto chromosome 2M^b^

Genome sequences of wheat landrace CS were used as references in Blastn searches to obtain positional information for unigenes annotated as R genes from *Ae. biuncialis* chromosome 2M^b^. Based on the chromosomal locations on CS Ref Seq v1.0, three comparative gene locus maps for the sequences of candidate disease resistance genes on 2M^b^ were made using MapDraw software referencing the locations of wheat homologous chromosome 2A, 2B and 2D.

### Results

### Cytogenetic analysis of CS-*Ae. biuncialis* 2M^b^ disomic addition line TA7733

Genomic *in situ* hybridization (GISH) and non-denaturing fluorescence *in situ* hybridization (ND-FISH) were respectively performed to confirm the chromosomal composition of CS-*Ae. biuncialis* 2M^b^ disomic addition line TA7733 by using fluorescein-labeled genomic DNA from M genome donor *Ae. comosa* as a probe and wheat CS DNA as blocker. As shown in Fig 1, there were 44 chromosomes including 42 wheat chromosomes and plus two *Ae. biuncialis* 2M^b^ chromosomes in TA7733.

**Fig 1.**
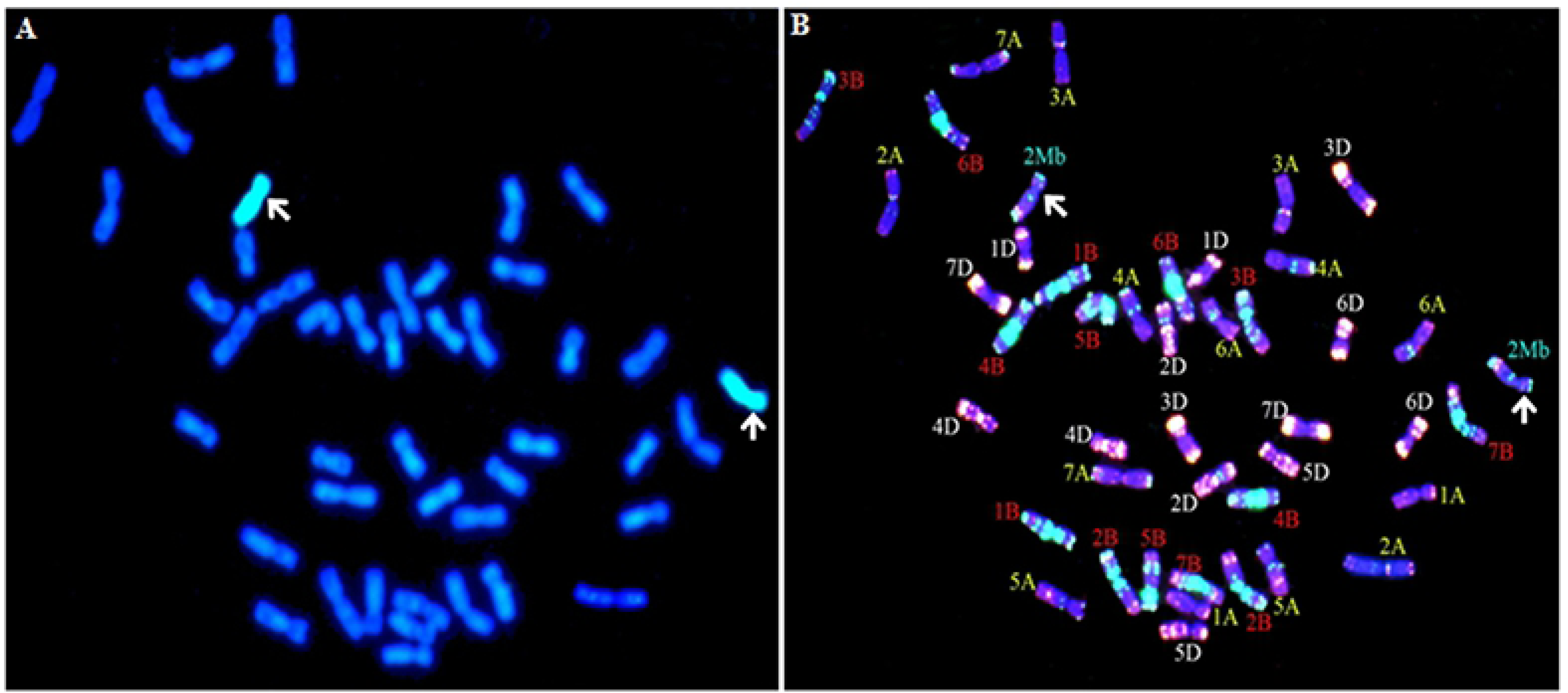
GISH and ND-FISH identification of CS-*Ae. biuncialis* 2M^b^ disomic addition line TA7733. (A) GISH patterns of CS-*Ae. biuncialis* 2M^b^ disomic addition line TA7733. Total genomic DNA of *Ae. comosa* was labelled with fluorescein-12-dUTP and visualized with green fluorescence. (B) ND-FISH patterns of CS-*Ae. biuncialis* 2M^b^ disomic addition line TA7733. Blue color indicated chromosomes counterstained with DAPI. Red color showed signals from oligos pAs1-3, pAs1-4, pAs1-6, AFA-3, AFA-4 and (AAC)10. Green color showed signals from oligos pSc119.2-1 and (GAA)_10_. The arrows indicated *Ae. biuncialis* chromosome 2M^b^.

### Assay of powdery mildew resistance of CS-*Ae. biuncialis* 2M^b^ disomic addition line TA7733

Prevalent *Bgt* isolates collected in Henan Province was used to inoculate seedlings with fully-extended first leaves of TA7733 and its recipient parent CS in the greenhouse. Ten days later, the leaves of CS covered with a large number of spores of powdery mildew, with infect types (IT) of 3-4, whereas TA7733 showed no spores, with ITs 0 to 1 (Fig 2A). Coomassie brilliant blue stained first leaf segments showed that CS was covered with hypha and spores had formed, while TA7733 only had a few blue spots (Fig 2B), confirming that TA7733 was high resistance to powdery mildew. Therefore the gene(s) conferring resistance to powdery mildew in TA7733 was mapped to chromosome 2M^b^ derived from *Ae. biuncialis.*

**Fig 2.**
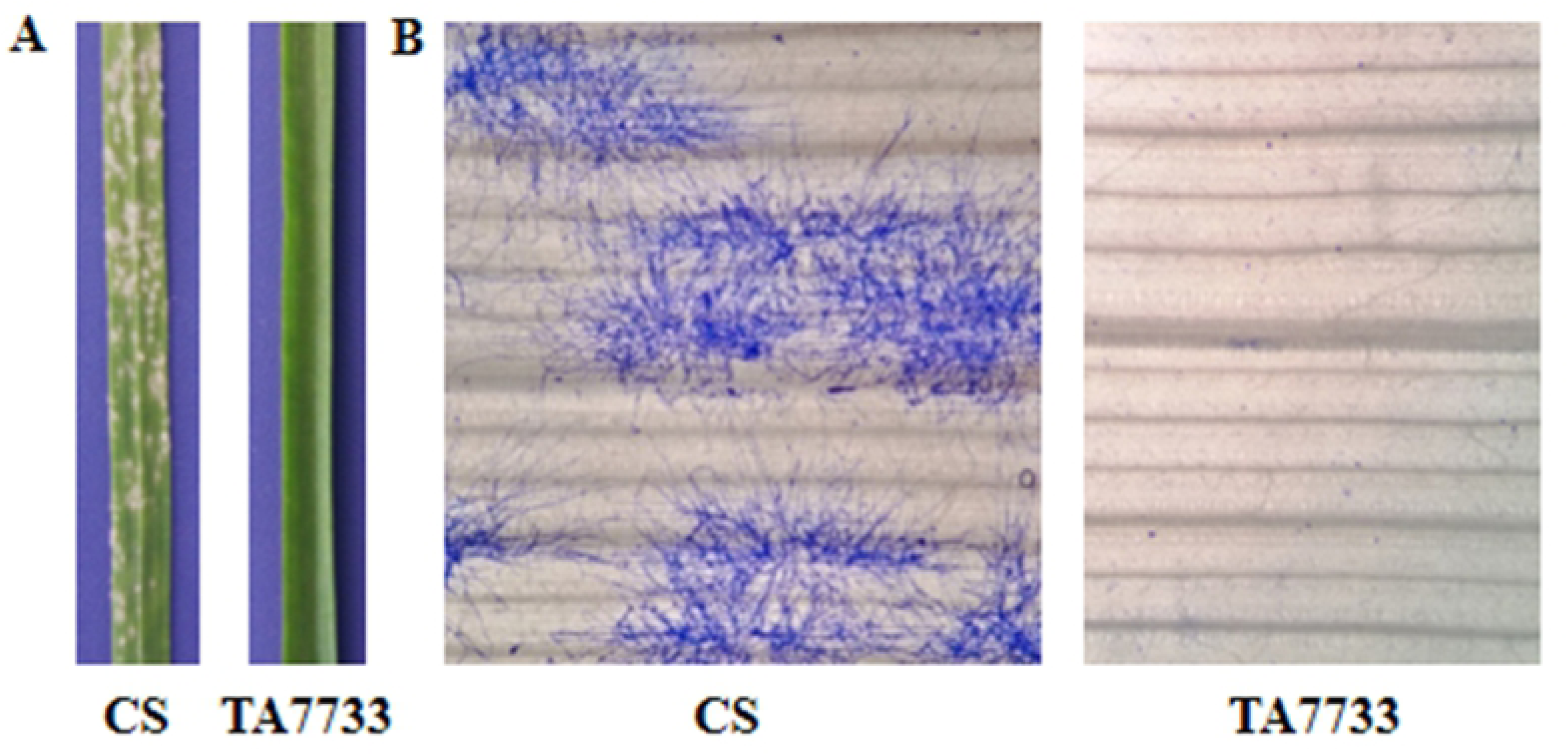
Powdery mildew resistance identification of CS-*Ae. biuncialis* 2M^b^ disomic addition line TA7733 and CS. (A) Disease symptoms of the first leaf of TA7733 and CS after 10 days of *Bgt* inoculation. (B) Microscopic analysis of the disease symptoms of TA7733 and CS by staining with coomassie brilliant blue-R-250.

Resistance spectrum of TA7733 was further assayed at seedling stage by inoculation of 15 prevalent *Bgt* isolates collected from different regions of China. As shown in Table 1, CS-*Ae. biuncialis* 2M^b^ disomic addition line TA7733 showed high resistance (IT = 0 or 1) to all the 15 *Bgt* isolates tested, whereas its recipient parent CS was highly susceptible (IT = 4 or 3) to all tested *Bgt* isolates. These results indicated that the gene(s) on chromosome 2M^b^ in TA7733 was a broad-spectrum resistance gene(s) to powdery mildew.

**Table 1.** Infection types of CS-*Ae. biuncialis* 2M^b^ disomic addition line TA7733 and CS for different *Bgt* isolates at the seedling stage.

| Isolates | Y01 | Y02 | Y03 | Y04 | Y05 | Y06 | Y07 | Y08 | Y09 | Y10 | Y11 | Y14 | Y15 | Y17 | Y18 |
| --- | --- | --- | --- | --- | --- | --- | --- | --- | --- | --- | --- | --- | --- | --- | --- |
| TA7733 | 0 | 0 | 0 | 0 | 0 | 1 | 0 | 0 | 0 | 0 | 1 | 0 | 0 | 0 | 0 |
| CS | 4 | 4 | 3 | 4 | 4 | 4 | 4 | 4 | 4 | 4 | 4 | 4 | 4 | 4 | 3 |

0 as immune, 0; as nearly immune, 1 as highly resistant, 2 as moderately resistant, 3 as moderately susceptible, 4 as highly susceptible.

### Transcriptome sequencing, *de novo* assembly and functional annotation

In order to discover resistance gene candidates and develop 2M^b^-specific molecular markers to assist the transfer resistance gene(s) at chromosome 2M^b^, RNA-seq of CS-*Ae. biuncialis* 2M^b^ disomic addition line TA7733 and its recipient parent CS were conducted before and after *Bgt*-infection. A total of 158,953 unigenes were assembled with a total length of 198,364,757 bp. Average unigene size was 1247.95 bp ranging from 301 to 19,496 bp. There were 89,951 unigenes (56.59%) in the length range of 301-1,000 bp, 42,478 unigenes (26.72%) in the length range of 1,001-2,000 bp, and 26,524 unigenes (16.69%) with length > 2,000 bp (Fig 3). All the unigenes were annotated with Blastx to the six public databases (NCBI NR protein, SWISSPROT, GO, KOG, KEGG and PRG databases) using a cutoff E-value of 10^-5^, giving rise to 86,196 (54.23%), 48,724 (30.65%), 40,543 (25.51%), 37,008 (23.28%), 13,414 (8.44%) and 10,969 (6.92%) annotated unigenes, respectively (Table 2). And 86,862 (54.65%) unigenes matched to at least one of the databases.

**Table 2.** Functional annotation of the unigenes by transcriptome sequencing of TA7733 during *Bgt* infection.

| database | NR | SWISSPROT | KOG | KEGG | GO | PRG | anno-union |
| --- | --- | --- | --- | --- | --- | --- | --- |
| annotation numbers | 86196 | 48724 | 37008 | 13414 | 40543 | 10969 | 86862 |
| annotation ratio (%) | 54.23 | 30.65 | 23.28 | 8.44 | 25.51 | 6.90 | 54.65 |

**Fig 3.**
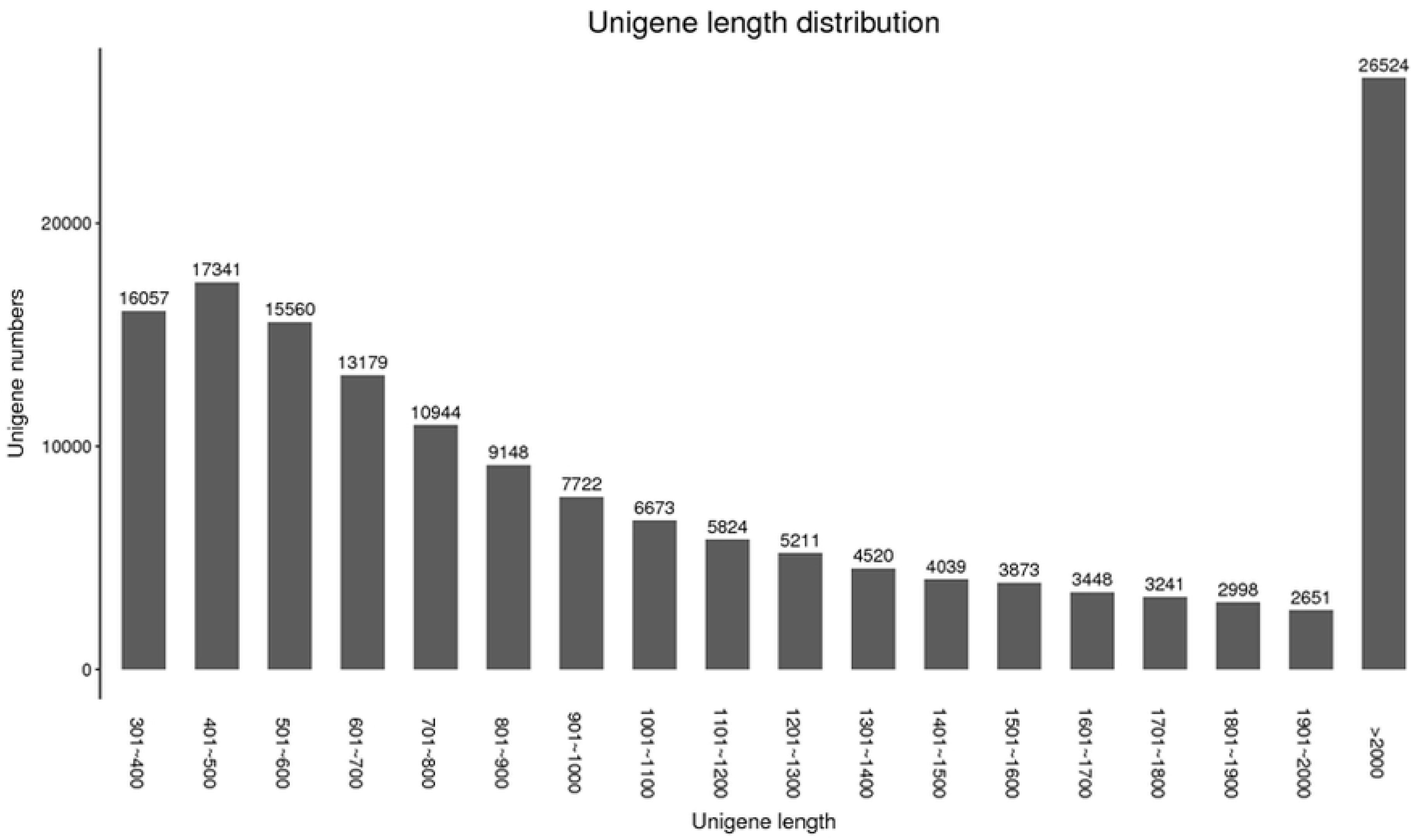
Length distribution of the assembled transcripts of CS-*Ae. biuncialis* 2M^b^ disomic addition line TA7733.

GO is an international classification system for standardized gene functions, which have three categories: biological process, molecular function and cellular component. According to sequence homology, a total of 40,543 (25.51% of 158,953) unigenes were assigned to one or more GO term annotations (Fig 4, S1 Table). Among which, “cellular process” (27,404; 67.59% of 40,543), “metabolic process” (24,470; 60.35% of 40,543), and “single-organism process” (20,606; 50.82% of 40,543) were the cardinal terms in the biological process category. “Cell” (30,742; 75.82% of 40,543), “cell part” (30,695; 75.71% of 40,543), and “organelle” (24,100; 59.44% of 40,543) were the most abundant terms in the cellular component category. “Binding” (24,379; 60.13% of 40,543) and “catalytic activity” (21,879; 53.96% of 40,543) were the most representative terms in the molecular function category. Only a few unigenes assigned into the terms of “extracellular matrix part” (9; 0.02% of 40,543), “protein tag” (8; 0.02% of 40,543) and “receptor regulator activity” (1; 0.00024% of 40,543).

**Fig 4.**
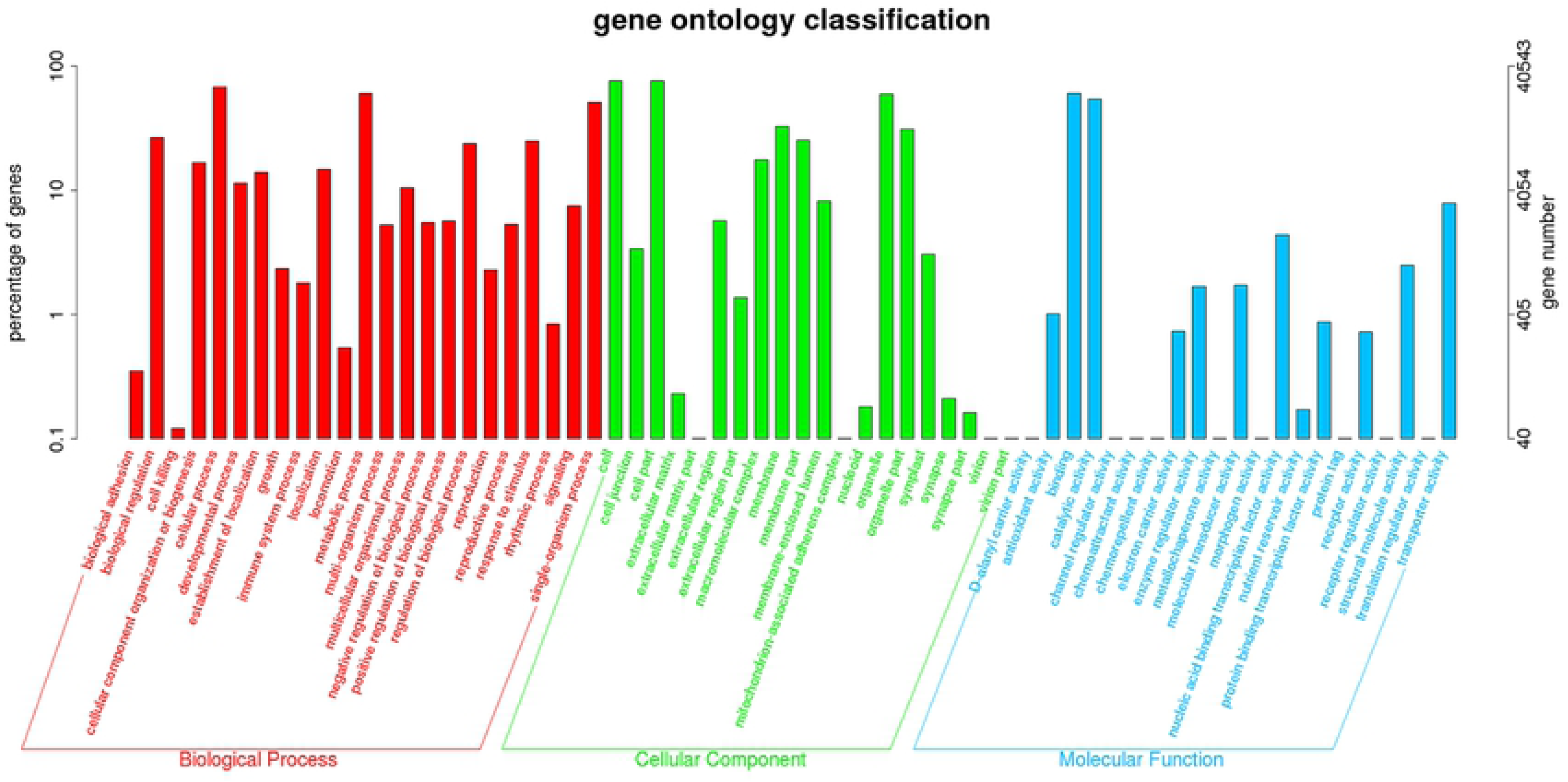
Histogram of GO categories of unigenes of CS-*Ae. biuncialis* 2M^b^ disomic addition line TA7733.

The KEGG database was used to systematically describe the pathway annotation of the unigenes. Out of total 158,953 annotated unigenes, 26,589 unigenes were assigned to 23 KEGG pathways (Fig 5, S2 Table). The most representative pathways of unigenes were the metabolic pathways (11920, 44.83%), genetic information processing (5456, 20.52%), environmental information processing (5095, 19.16%) and cellular processes (4118, 15.49%).

**Fig 5.**
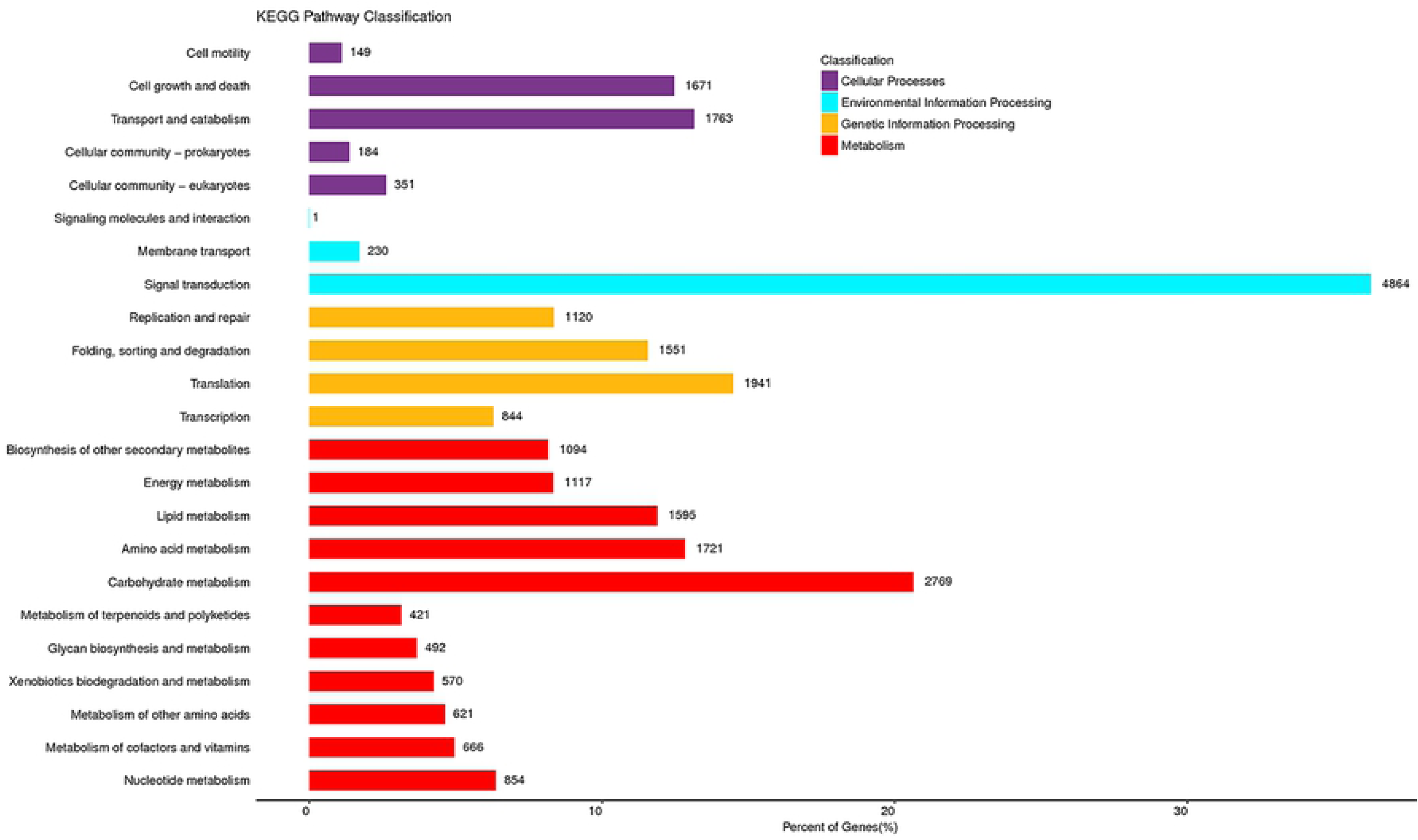
Clusters of KEGG functional classifications of unigenes of CS-*Ae. biuncialis* 2M^b^ disomic addition line TA7733.

### Analyses of genes involved in responses to *Bgt* infection from *Ae. biuncialis* chromosome 2M^b^

As many as 158,953 unigenes involved in responses to *Bgt* infection have been identified from TA7733. In this study, we aimed to explore the resistance-related genes derived from *Ae. biuncialis* chromosome 2M^b^. So we primarily focused on the genes specifically expressed in TA7733 which were not expressed in CS. These genes were most likely derived from *Ae. biuncialis* chromosome 2M^b^.

Among the total number of 158,953 unigenes, there were 7,278 unigenes which were expressed in TA7733 but not in CS before and after *Bgt*-inoculation. Of which, 4,382 unigenes were differentially expressed genes (DEGs) in TA7733 post vs before *Bgt*-inoculation, and the remaining 2,896 unigenes had expression levels of no significant difference. In consideration of the fact that expression levels of some cloned resistance genes did show no significant difference before and after pathogen infection [9,41], we also take into account these 2,896 unigenes as putative candidate genes involved in powdery mildew resistance from chromosome 2M^b^.

To analyze the biological pathways involved in resistance after inoculation with *Bgt*, the KOBAS software was used to test the statistical enrichment of DEGs in KEGG pathways and it was found 399 out of 4,382 DEGs were allocated to 162 KEGG pathways (S3 Table). The most representative pathways were the phenylpropanoid biosynthesis (28, 7.02%), plant hormone signal transduction (23, 5.76%), flavonoid biosynthesis (15, 3.76%), stilbenoid, diarylheptanoid and gingerol biosynthesis (14, 3.51%) and MAPK signaling pathway-plant (14, 3.51%), followed by glutathione metabolism (13, 3.26%), protein processing in endoplasmic reticulum (13, 3.26%), and metabolism of xenobiotics by cytochrome P450 (12, 3.00%). These annotations provided valuable clues in investigation of the specific processes and identification of the genes that involved in powdery mildew resistance conferred by *Ae. biuncialis* 2M^b^ chromosome.

### Selection and verification of disease resistance gene candidates of chromosome 2M^b^

Based on transcriptome data analysis, 7,278 unigenes specific to *Ae. biuncialis* chromosome 2M^b^ were checked out. Among these 7,278 unigenes, 295 unigenes were annotated as R genes by Blastx alignment against the PRG and NCBI databases. Sequences of these 295 R genes were used to blastn against with CS Ref Seq v1.0 to map these genes to wheat homologous groups. As a result, sixty-one (61/295, 20.68%) R genes were mapped to wheat homologous group 2.

In order to verify if these 61 R genes were derived from *Ae. biuncialis* chromosome 2M^b^, a total of 61 sets of PCR primers were designed based on their transcriptome sequences. PCR amplification of genomic DNA of CS and TA7733 confirmed that 40 R genes were specific to chromosome 2M^b^ in CS-*Ae. biuncialis* 2M^b^ disomic addition line TA7733 (Fig 6, S4 Table). So these 40 R genes could be considered as candidates involved in powdery mildew resistance derived from *Ae. biuncialis* 2M^b^ chromosome.

**Fig 6.**
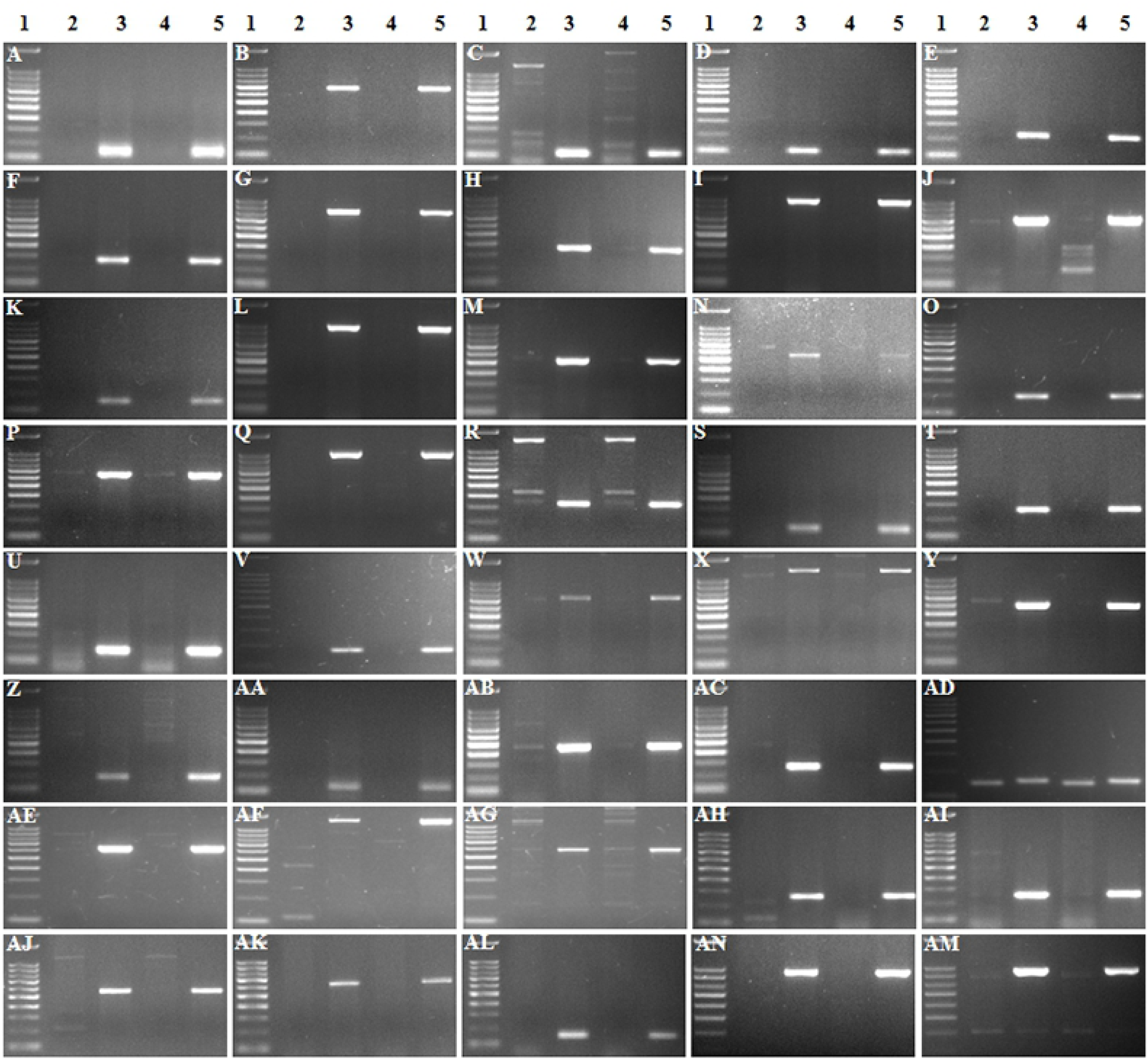
PCR amplification patterns of 40 *Ae. biuncialis* chromosome 2M^b^-specific molecular markers. (1) 100 bp DNA Marker. (2, 4) CS. (3, 5) TA7733. (A) CL119404Contig1. (B) CL88277Contig1. (C) CL82670Contig1. (D) 82789Contig1. (E) CL82700Contig1. (F) CL85355Contig1. (G) CL66003Contig1. (H) CL89405Contig1. (I) CL106750Contig1. (J) CL52922Contig1. (K) CL19981Contig2. (L) CL93721Contig1. (M) CL84424Contig1. (N) CL88613Contig1. (O) CL91022Contig1. (P) 96221Contig1. (Q) comp19533_c0_seq1_6. (R) CL85258Contig1. (S) CL113849Contig1. (T) CL86521Contig1. (U) CL105879Contig1. (V) CL90029Contig1. (W) CL84846Contig2. (X) CL87530Contig1. (Y) CL29910Contig1. (Z) CL92547Contig1. (AA) CL75219Contig1. (AB) CL108886Contig1. (AC) CL113652Contig1. (AD) CL80063Contig1. (AE) CL89447Contig1. (AF) CL93169Contig1. (AG) CL114224Contig1. (AH) CL116612Contig1. (AI) CL67241Contig1. (AJ) CL119539Contig1. (AK) CL90483Contig1. (AL) CL91742Contig1. (AM) comp84147_c0_seq1_6. (AN) CL124Contig7.

Among the 40 R genes (Tables 3 and 4, S5 and S6 Tables), 21 genes were assigned by alignment to PRG database. Of these 21 R genes, 11 R genes were in CNL class containing a predicted coiled-coil (CC) structures, a central nucleotide-binding (NB) subdomain and a leucine rich repeat (LRR) domain, six in NL class containing NBS and LRR domains, but lack of CC domain, three in class RLP which contains leucine-rich receptor-like repeat, a transmembrane region of 25AA, and a short cytoplasmic region, and one in TNL class which contains a central NB subdomain, a LRR domain, a interleukin-1 receptor (1L-1R) domain (Table 4). The remaining 19 R genes were aligned to NCBI database, encoding protein kinase, disease resistance protein RGA, disease resistance protein RP and cytochrome P450, respectively.

**Table 3.** The functions of 40 disease resistance gene candidates from *Ae. biuncialis* chromosome 2M^b^.

| Unigene IDs | Gene annotation | up-down | Similarity with wheat homoeologous group 2 |
| --- | --- | --- | --- |
| **CL84424Contig1<sup>a</sup>** | Cysteine-rich receptor-like protein kinase 26 [*Ae. tauschii*] | unregulated | 0.83 |
| **CL93721Contig1<sup>a</sup>** | G-type lectin S-receptor-like serine/threonine-protein kinase At1g11300 [*Hordeum vulgare*] | unregulated | 0.90 |
| **CL89447Contig1<sup>b</sup>** | LRR and NB-ARC domains-containing disease resistance protein (best arabidopsis hit); NBS-LRR disease resistance protein, putative, expressed (best rice hit) (CNL) | unregulated | 0.82 |
| **CL90029Contig1<sup>b</sup>** | NBS-LRR disease resistance protein-like protein (NBS-LRR1) [*H. vulgare*] (CNL) | unregulated | 0.88 |
| **CL88613Contig1<sup>a</sup>** | Predicted: disease resistance protein RGA2-like [*Brachypodium distachyon*] | unregulated | 0.86 |
| **CL96221Contig1<sup>a</sup>** | Cysteine-rich receptor-like protein kinase 10 [*T. urartu*] | unregulated | 0.79 |
| **CL106750Contig1<sup>b</sup>** | Putative disease resistance protein RGA1 [*Ae. tauschii*] (CNL) | unregulated | 0.82 |
| **CL113949Contig1<sup>a</sup>** | Putative disease resistance protein RGA1 [*Ae. tauschii*] | unregulated | 0.78 |
| **CL91742Contig1<sup>b</sup>** | (CNL) | unregulated | 0.78 |
| **CL116612Contig1<sup>b</sup>** | Putative disease resistance RPP13-like protein 1 [*T. urartu*] (CNL) | unregulated | 0.89 |
| **CL82670Contig1<sup>a</sup>** | Cytochrome P450 71D7 [*Ae. tauschii*] | unregulated | 0.72 |
| **CL93169Contig1<sup>b</sup>** | NB-ARC domain-containing disease resistance protein (best arabidopsis hit); RGH1A, putative, expressed (best rice hit) (NL) | unregulated | 0.79 |
| **CL108886Contig1<sup>b</sup>** | LRR and NB-ARC domains-containing disease resistance protein (best arabidopsis hit); NBS-LRR disease resistance protein, putative, expressed (best rice hit) (CNL) | unregulated | 0.85 |
| **comp19533_c0_seq1_6<sup>b</sup>** | LRR and NB-ARC domains-containing disease resistance protein (best arabidopsis hit); NBS-LRR disease resistance protein, putative, expressed (best rice hit) (CNL) | unregulated | 0.88 |
| **CL90483Contig1<sup>b</sup>** | Putative disease resistance RPP13-like protein 1 [*T. urartu*] (NL) | unregulated | 0.83 |
| **CL85355Contig1<sup>a</sup>** | Disease resistance protein RPM1 [*T. urartu*] | unregulated | 0.80 |
| **CL80063Contig1<sup>b</sup>** | Leucine-rich repeat protein kinase family protein (best arabidopsis hit) (RLP) | unregulated | 0.80 |
| **CL66003Contig1<sup>b</sup>** | Putative disease resistance protein RGA4 [*Ae. tauschii*] (NL) | unregulated | 0.76 |
| **CL119404Contig1<sup>b</sup>** | Putative disease resistance RPP13-like protein 1 [*Ae. tauschii*] (CNL) | unregulated | 0.88 |
| **CL113652Contig1<sup>b</sup>** | Predicted: putative disease resistance RPP13-like protein 3 (LOC109731753) [*Ae. tauschii*] (NL) | unregulated | 0.88 |
| **CL91022Contig1<sup>a</sup>** | Lectin-domain containing receptor kinase A4.3 [*Ae. tauschii*] | unregulated | 0.78 |
| **CL85258Contig1<sup>a</sup>** | Predicted: G-type lectin S-receptor-like serine/threonine-protein kinase B120 [*Brachypodium distachyon*] | unregulated | 0.77 |
| **CL105879Contig1<sup>b</sup>** | Putative LRR receptor-like serine/threonine-protein kinase [*Ae. tauschii*] (RLP) | unregulated | 0.89 |
| **CL84846Contig1<sup>a</sup>** | Cysteine-rich receptor-like protein kinase 29 [*Ae. tauschii*] | unregulated | 0.90 |
| **CL67241Contig1<sup>b</sup>** | (CNL) | unregulated | 0.77 |
| **CL124Contig7<sup>b</sup>** | HOPZ-ACTIVATED RESISTANCE 1 (best arabidopsis hit); Leucine Rich Repeat family protein, expressed (best rice hit) (NL) | unregulated | 0.85 |
| **CL89405Contig1<sup>a</sup>** | Putative disease resistance protein RGA4 [*Ae. tauschii*] | unregulated | 0.86 |
| **CL119216Contig1<sup>a</sup>** | Putative serine/threonine-protein kinase receptor [*Ae. tauschii*] | unregulated | 0.86 |
| **CL119539Contig1<sup>b</sup>** | (TNL) | unregulated | 0.88 |
| **CL86521Contig1<sup>a</sup>** | Putative serine/threonine-protein kinase-like protein CCR3 [*Ae. tauschii*] | unregulated | 0.83 |
| **CL29910Contig1<sup>b</sup>** | Disease resistance protein RGA2 [*Ae. tauschii*] (NL) | unregulated | 0.68 |
| **CL87530Contig1<sup>a</sup>** | Wall-associated receptor kinase 4 [*T. urartu*] | unregulated | 0.78 |
| **CL114224Contig1<sup>b</sup>** | NB-ARC domain-containing disease resistance protein (best arabidopsis hit); NB-ARC domain containing protein, expressed (best rice hit) (CNL) | unregulated | 0.88 |
| **CL82700Contig1<sup>a</sup>** | Lectin-domain containing receptor kinase A4.3 [*Ae. tauschii*] | unregulated | 0.80 |
| **comp84147_c0_seq1_6<sup>b</sup>** | TSA: Triticum aestivum cultivar Bobwhite isotig02189.flagleaf mRNA sequence (CNL) | upregulated | 0.88 |
| **CL92547Contig1<sup>a</sup>** | Predicted: probable LRR receptor-like serine/threonine-protein kinase At1g05700 (LOC109742478) [*Ae. tauschii*] | upregulated | 0.87 |
| **CL82789Contig1<sup>a</sup>** | Putative LRR receptor-like serine/threonine-protein kinase [*Ae. tauschii*] | upregulated | 0.77 |
| **CL88277Contig1<sup>b</sup>** | Predicted: probable leucine-rich repeat receptor-like protein kinase At1g35710 (LOC109774313) [*Ae. tauschii*] (RLP) | upregulated | 0.85 |
| **CL19981Contig2<sup>a</sup>** | Putative disease resistance protein RGA3 [*Ae. tauschii*] | downregulated | 0.80 |
| **CL75219Contig1<sup>a</sup>** | Predicted: putative disease resistance RPP13-like protein 3 (LOC109732887) [*Ae. tauschii*] | downregulated | 0.89 |

<sup>a</sup> indicated these R genes were assigned by alignment to NCBI database. <sup>b</sup> indicated these R genes were assigned by alignment to PRG database.

**Table 4.**
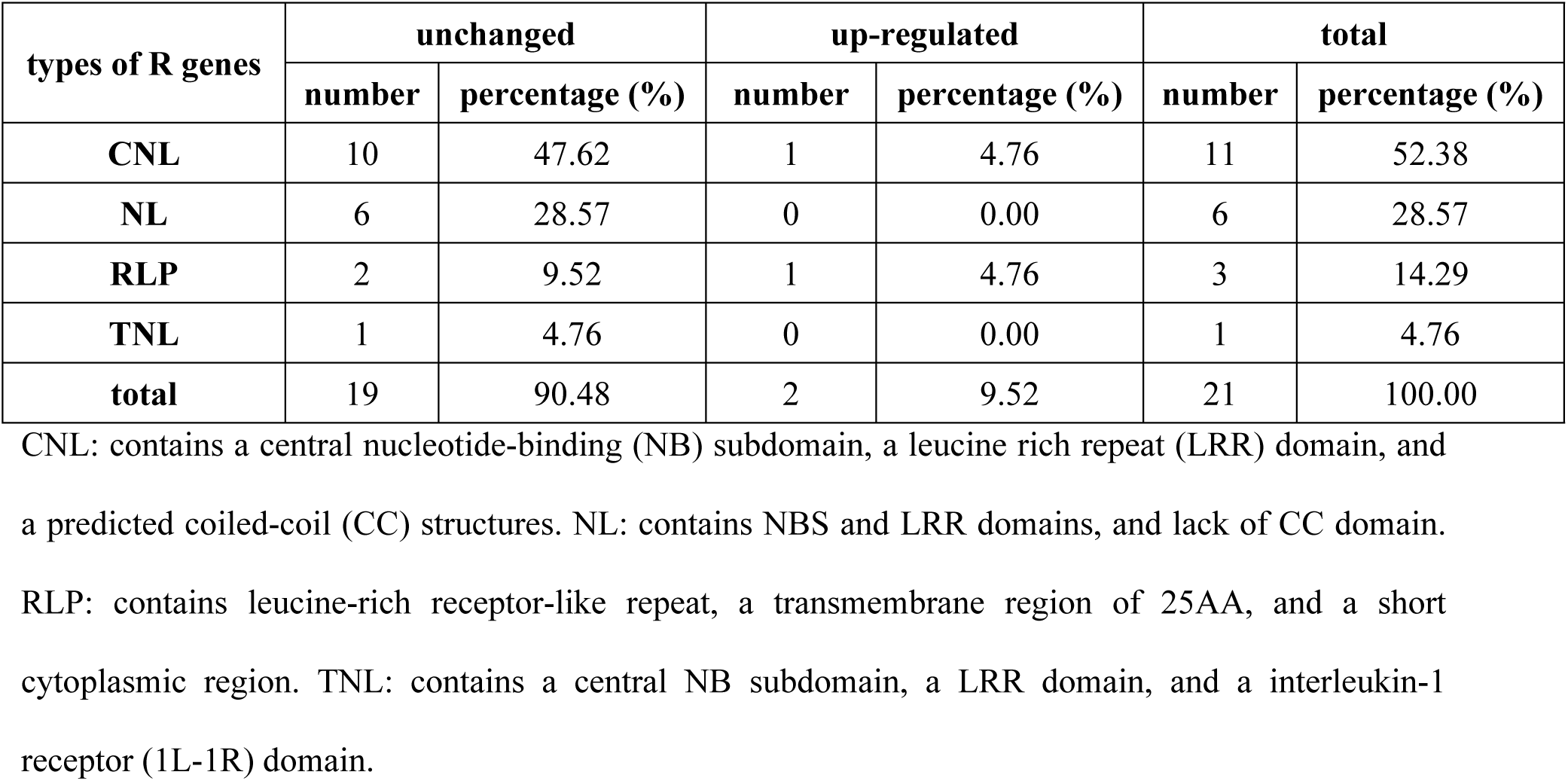
The types of unigenes annotated as R genes alignment against the PRG database from *Ae. biuncialis* chromosome 2M^b^ during *Bgt* infection.

Comparative mapping was carried out by using MapDraw software based on alignment of sequences of these 40 R genes of *Ae. biuncialis* chromosome 2M^b^ specific with those in CS Ref Seq v1.0 (Fig 7). The maps showed that there were 21 R genes were assigned to terminal of the long arm, 16 to the terminal of the short arm of chromosome 2M^b^, respectively. Besides, two R genes (CL75219contig1 and CL92547contig1) mapped to the location close to the centromere of long arm, and one R genes (CL96221contig1) neighbored to the centromere of short arm of chromosome 2M^b^.

**Fig 7.**
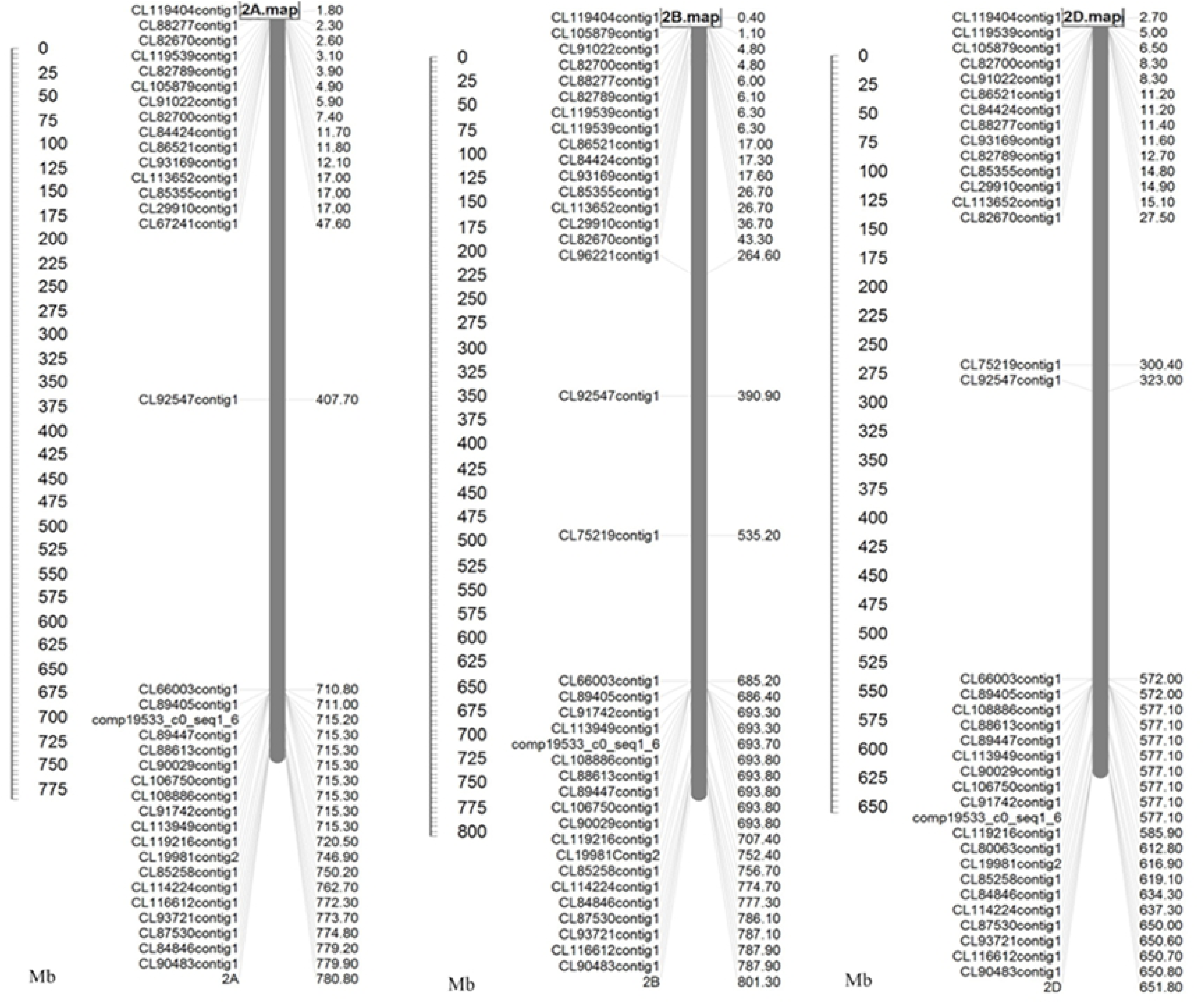
Comparative mapping of 40 2M^b^-specific R genes by alignment with CS Ref Seq v1.0.

## Discussion

Development of resistant wheat varieties by utilization of resistance genes is the most important and environment-friendly way to control *Bgt-*caused damages. The genes with broad spectrum and durability resistance make it highly valuable in wheat breeding programs [29]. Wild relatives of common wheat harbored considerable genetic diversity for powdery mildew resistance. For example, the wild relatives of common wheat, *Secale cereale, Dasypyrum villosum* and *Ae. searsii* conferred powdery mildew resistance gene *Pm7, PmJZHM2RL, Pm62* and *Pm57* from homoeologous group 2 [38, 42-44]. In present study, resistance spectrum of the CS-*Ae. biuncialis* 2M^b^ disomic addition line TA7733 was assayed by using 15 *Bgt* isolates originating from different wheat-producing regions in China. The results showed TA7733 were resistance to all the isolates, indicating that the gene(s) on *Ae. biuncialis* 2M^b^ chromosome was broad-spectrum powdery mildew resistance. Currently no any other catalogued powdery mildew resistance genes were reported to be derived from *Ae. biuncialis* homoeologous group 2. So the gene(s) on *Ae. biuncialis* 2M^b^ chromosome should be a novel resistance gene(s) to wheat powdery mildew.

Previous studies have generally focused on the significantly differentially expressed genes in interactions between plant and pathogens to discover disease resistance-related genes by transcriptome sequencing [45-47]. However, some cloned genes of disease resistance in plants were reported to be not significantly up-regulated expression after pathogens inoculation. For example, Zou et al. (2017) cloned wheat powdery mildew resistance gene *Pm60* using standard map-based cloning, and showed that the transcript levels of *Pm60* were no significantly difference after *Bgt* E09 infection at various time points by qRT-PCR [9]. In addition, Li et al. (2017) discovered the expression level of broad-spectrum blast resistance gene *bsr-d1* in rice was also not induced before and after blast infection [41]. So, the chance to obtain candidate disease genes might be lost if only significantly regulated genes were considered as R gene candidates. Therefore, in this study we explored *Bgt*-resistance related candidate genes from all unigenes of 2M^b^-specific regardless they were significantly regulated or not induced post *Bgt* infection. After PCR verification of chromosome 2M^b^ specificity by using R gene-based primer sets, we finally selected 40 R genes candidates including 34 unregulated, four up-regulated and two down-regulated genes.

Isolation of plant resistance (R) gene is greatly helpful to breed resistant varieties and elucidates resistance molecular mechanisms. Conventional map-based cloning proved to be a effective method to clone R genes [45]. It is time consuming and difficult to fine map R genes from wild relatives of common wheat due to lack of exchange and combination between the alien chromatin and wheat homoeologous counterpart. In previous study, more than 70 R genes against diverse pathogens have been isolated from various plant through map-based cloning [48]. And nearly three quarters of R genes cloned so far encoded NBS-LRR protein [48]. NBS-LRR R genes reportedly recognized pathogens and initiated defense responses subsequently. To date, five out of 89 *Pm* genes identified in wheat gene pool, such as *Pm2* [49], *Pm3* [50], *Pm8* [51], *Pm21* [28,52] and *Pm60* [9], have been cloned, which all encoded CC-NBS-LRR proteins. So CC-NBS-LRR genes identification could provide a promising strategy to isolate candidate R genes. In this study, 11 out of 40 2M^b^-specific R genes were predicted to encode CC-NBS-LRR protein. These 11 R genes may be considered as the most important candidates for further isolating and cloning R genes from *Ae. biuncialis* chromosome 2M^b^.

GISH is a popular visual method to identify alien chromosome or chromatin in wheat background. Whereas GISH is expensive and time-consuming especially used to screen a large population derived from distant crossing between wheat and its wild relatives [53]. In contrast, molecular markers are not affected by environmental conditions, tissue or developmental stage and gene expression, and possess high genetic polymorphism [54,55]. So the development and application of molecular markers have been considered as new and low-cost ways to quickly identify alien chromosomes or chromatin. High-throughput RNA-seq technology can generate large amounts of transcriptome sequences and has been widely used to develop molecular markers specific to chromosomes of wild relatives of wheat, especially those with limited public genomic sequences. Li et al. (2017) developed 25 *D. villosum* 6V#4S-specifc markers using transcriptome data [56]. Wang et al. (2018) developed 134 *Ae. longissima* chromosome*-*specific markers by RNA-seq [30]. Li et al. (2019) developed 76 molecular markers specific to the chromosome 1V to 7V of *D. villosum*#4 based on transcriptome data [31]. The transcription sequences were highly conserved and might be associated with the genes that related to a definite trait [57]. The markers developed by transcriptome sequencing will help to identify candidate functional genes and increase the efficiency of marker-assisted selection [57,58]. In this study, 40 functional molecular markers of R genes specific to *Ae. biuncialis* chromosome 2M^b^, including 23 and 17 markers located on the long and short arm of chromosome 2M^b^, respectively, were developed by transcriptome data analyses. These markers will be useful to assist selection of resistant wheat-*Ae. biuncialis* 2M^b^ translocations and transfer resistance gene(s) from 2M^b^ to common wheat by inducing CS-*Ae. biuncialis* homoeologous recombination for wheat disease breeding in the future.

## Conclusions

In summary, powdery mildew resistance gene(s) on *Ae. biuncialis* chromosome 2M^b^ was verified to be board-spectrum in this study. It could be a valuable disease-resistance resource for wheat breeding programs. Forty candidate disease resistance genes selected based on transcriptome sequencing analyses will be greatly helpful to isolate and clone disease resistance gene(s) derived from chromosome 2M^b^ and provide the insight into the molecular mechanism of disease resistance. Furthermore, 40 functional molecular markers of those R gene candidates of 2M^b^ specific in this study would facilitate transferring resistance gene(s) from 2M^b^ to common wheat by inducing CS-*Ae. biuncialis* homoeologous recombination.

## Acknowledgments

We sincerely thank Dr. Yuli Song from the Institute of Plant Protection, Henan Academy of Agricultural Sciences, Henan Province, China for providing prevailing *Bgt* isolates collected in Henan.

## Supporting information

**S1 Table. Gene ontology of transcriptome of CS-*Ae. biuncialis* 2M**^**b**^ **disomic addition line TA7733.**

**S2 Table. The KEGG pathway classification of transcriptome of CS-*Ae. biuncialis* 2M**^**b**^ **disomic addition line TA7733.**

**S3 Table. KEGG pathway classification of DEGs of CS-*Ae. biuncialis* 2M**^**b**^ **disomic addition line TA7733.**

**S4 Table. The 40 *Ae. biuncialis* chromosome 2M**^**b**^**-specifc markers developed in this study based on unigenes annotated as R genes.**

**S5 Table. The expression levels of 40 candidate disease resistance genes from *Ae. biuncialis* chromosome 2M**^**b**^.

**S6 Table. The list sequences of 40 candidate disease resistance genes from *Ae. biuncialis* chromosome 2M**^**b**^.

## References

1. Ramirez-Gonzalez RH, Segovia V, Bird N, Fenwick P, Holdgate S, Berry S, et al. RNA-Seq bulked segregant analysis enables the identification of high-resolution genetic markers for breeding in hexaploid wheat. Plant Biotechnol J. 2015; 13: 613–624.

2. Fu DL, Uauy C, Distelfeld A, Blechl A, Epstein L, Chen XM, et al. A kinase-START gene confers temperature-dependent resistance to wheat stripe rust. Science. 2009; 323: 1357–1360.

3. Griffey C, Das M, Stromberg EJ. Effectiveness of adult-plant resistance in reducing grain yield loss to powdery mildew in winter wheat. Plant Dis. 1993; 77: 618–622.

4. Li GQ, Carver BF, Cowger C, Bai GH, Xu XY. *Pm223899*, a new recessive powdery mildew resistance gene identified in Afghanistan landrace PI223899. Theor Appl Genet. 2018; 131: 2775–2783.

5. Ma PT, Xu HX, Luo QL, Qie YM, Zhou YL, Xu YF, et al. Inheritance and genetic mapping of a gene for seedling resistance to powdery mildew in wheat line X3986-2. Euphytica. 2014; 200: 149–157.

6. Morgounov A, Tufan HA, Sharma R, Akin B, Bagci A, Braun HJ, et al. Global incidence of wheat rusts and powdery mildew during 1969-2010 and durability of resistance of winter wheat variety Bezostaya 1. Eur J Plant Pathol. 2012; 132: 323–340.

7. Huang J, Zhao ZH, Song FJ, Wang XM, Xu HX, Huang Y, et al. Molecular detection of a gene effective against powdery mildew in the wheat cultivar Liangxing 66. Mol Breed. 2012; 30: 1737–1745.

8. Zhang DY, Zhu KY, Dong LL, Liang Y, Li GQ, Fang TL, et al. Wheat powdery mildew resistance gene Pm64 derived from wild emmer (Triticum turgidum var. dicoccoides) is tightly linked in repulsion with stripe rust resistance gene Yr5. 2019; https://doi.org/10.1016/j.cj.2019.03.003.

9. Zou SH, Wang H, Li YW, Kong ZS, Tang DZ. The NB-LRR gene *Pm60* confers powdery mildew resistance in wheat. New Phytol. 2018; 218: 298–309.

10. McDonald BA, Linde C. Pathogen population genetics, evolutionary potential, and durable resistance. Annu Rev Phytopathol. 2002; 40: 349–379.

11. Xiao MG, Song FJ, Jiao JF, Wang XM, Xu HX, Li HJ. Identification of the gene *Pm47* on chromosome 7BS conferring resistance to powdery mildew in the Chinese wheat landrace Hongyanglazi. Theor Appl Genet. 2013; 126: 1397–1403.

12. Ma PT, Xu HX, Zhang HX, Li LH, Xu YF, Zhang XT, et al. The gene *PmWFJ* is a new member of the complex *Pm2* locus conferring unique powdery mildew resistance in wheat breeding line Wanfengjian 34. Mol Breed. 2015; 35: 210.

13. Summers R, Brown J. Constraints on breeding for disease resistance in commercially competitive wheat cultivars. Plant Pathol. 2013; 62: 115–121.

14. Resta P, Zhang HB, Dubcovsky J, Dvořák J. The origins of the genomes of *Triticum biunciale, T. ovatum, T. neglectum, T. columnare*, and *T. rectum* (Poaceae) based on variation in repeated nucleotide sequences. Am J Bot. 1996; 83: 1556–1565.

15. Badaeva E, Amosova A, Samatadze T, Zoshchuk S, Shostak N, Chikida N, et al. Evolution, Genome differentiation in *Aegilops*. 4. Evolution of the U-genome cluster. Plant Sys Evol. 2004; 246: 45–76.

16. Damania AB, Pecetti L. Variability in a collection of *Aegilops* species and evaluation for yellow rust resistance at two locations in northern Syria. J Genet Breed. 1990; 44: 97–102.

17. Dimov A, Zaharieva M, Mihova S. Rust and powdery mildew resistance in Aegilops accessions from Bulgaria. In: Damania AB, editors. 1993. pp. 165–169.

18. Makkouk K, Ghulam W, Comeau A. Resistance to barley yellow dwarf luteovirus in *Aegilops* species. Can J Plant Sci. 1994; 74: 631–634.

19. Zhao H, Zhang W, Wang J, Li F, Cui F, Ji J, et al. Comparative study on drought tolerance of wheat and wheat-*Aegilops biuncialis* 6Ub addition lines. J Food Agric Environ. 2013; 11: 1046–1052.

20. Molnár I, Gáspár L, Sárvári É, Dulai S, Hoffmann B, Molnár-Láng M, et al. Physiological and morphological responses to water stress in *Aegilops biuncialis* and *Triticum aestivum* genotypes with differing tolerance to drought. Funct Plant Biol. 2004; 31: 1149–1159.

21. Colmer TD, Flowers TJ, Munns, R. Use of wild relatives to improve salt tolerance in wheat. J Exp Bot. 2006; 57: 1059–1078.

22. Farkas A, Molnár I, Dulai S, Rapi S, Oldal V, Cseh A, et al. Increased micronutrient content (Zn, Mn) in the 3M^b^(4B) wheat-*Aegilops biuncialis* substitution and 3Mb.4BS translocation identified by GISH and FISH. Genome. 2014; 57: 61–67.

23. Zhou JP, Yao CH, Yang EN, Yin MQ, Liu C, Ren ZL. Characterization of a new wheat-*Aegilops biuncialis* addition line conferring quality-associated HMW glutenin subunits. Genet Mol Res. 2014; 13: 660–669.

24. Schneider A, Linc G, Molnár I, Molnár-Láng M. Molecular cytogenetic characterization of *Aegilops biuncialis* and its use for the identification of 5 derived wheat-*Aegilops biuncialis* disomic addition lines. Genome. 2005; 48: 1070–1082.

25. Schneider A, Molnar-Lang M. Detection of various U and M chromosomes in wheat-*Aegilops biuncialis* hybrids and derivatives using fluorescence *in situ* hybridisation and molecular markers. Czech J Genet Plant Breed. 2012; 48: 169–177.

26. Xia Q, Mai YN, Dong ZJ, Liu WX. Identification of powdery mildew resistance resources from wheat-wild relative disomic addition lines and development of molecular markers of alien chromosome-specialty. J Henan Agric Sci. 2018; 47: 64–69.

27. Wang ZZ, Li HW, Zhang DY, Guo L, Chen JJ, Chen YX, et al. Genetic and physical mapping of powdery mildew resistance gene *MlHLT* in Chinese wheat landrace Hulutou. Theor Appl Genet. 2015; 128: 365–373.

28. He HG, Zhu SY, Zhao RH, Jiang ZN, Ji YY, Ji J, et al. *Pm21*, encoding a typical CC-NBS-LRR protein, confers broad-spectrum resistance to wheat powdery mildew disease. Mol Plant. 2018; 11: 879–882.

29. Cao AZ, Xing LP, Wang XY, Yang XM, Wang W, Sun YL, et al. Serine/threonine kinase gene *Stpk-V*, a key member of powdery mildew resistance gene *Pm21*, confers powdery mildew resistance in wheat. Proc Natl Acad Sci. 2011; 108: 7727–7732.

30. Wang KY, Lin ZS, Wang L, Wang K, Shi QH, D. LP, et al. Development of a set of PCR markers specific to *Aegilops longissima* chromosome arms and application in breeding a translocation line. Theor Appl Genet. 2018; 131: 13–25.

31. Li SJ, Wang J, Wang KY, Chen JN, Wang K, Du LP, et al. Development of PCR markers specific to *Dasypyrum villosum* genome based on transcriptome data and their application in breeding *Triticum aestivum*-*D. villosum*#4 alien chromosome lines. BMC Genomics. 2019; 20: 289.

32. Rubio M, Rodríguez-Moreno L, Ballester AR, Moura MC, Bonghi C, Candresse T, et al. Analysis of gene expression changes in peach leaves in response to *Plum pox virus* infection using RNA-Seq. Mol Plant Pathol. 2015; 16: 164–176.

33. Zhang H, Fu Y, Guo H, Zhang L, Wang CY, Song WN, et al. Transcriptome and proteome-based network analysis reveals a model of gene activation in wheat resistance to stripe rust. Int J Mol Sci. 2019; 20: 1106.

34. Zhang H, Yang YZ, Wang CY, Liu M, Li H, Fu Y, et al. Large-scale transcriptome comparison reveals distinct gene activations in wheat responding to stripe rust and powdery mildew. BMC genomics. 2014; 15: 898.

35. Li QQ, Niu ZB, Bao YG, Tian QJ, Wang HG, Kong LR, et al. Transcriptome analysis of genes related to resistance against powdery mildew in wheat-*Thinopyrum* alien addition disomic line germplasm SN6306. Gene. 2016; 590: 5–17.

36. Huang XY, Zhu MQ, Zhuang LF, Zhang SY, Wang JJ, Chen XJ, et al. Structural chromosome rearrangements and polymorphisms identified in Chinese wheat cultivars by high-resolution multiplex oligonucleotide FISH. Theor Appl Genet. 2018; 131: 1967–1986.

37. Li HH, Jiang B, Wang JC, Lu YQ, Zhang JP, Pan CL, et al. Mapping of novel powdery mildew resistance gene(s) from *Agropyron cristatum* chromosome 2P. Theor Appl Genet. 2017; 130: 109–121.

38. Liu WX, Koo DH, Xia Q, Li CX, Bai FQ. Song YL, et al. Homoeologous recombination-based transfer and molecular cytogenetic mapping of powdery mildew-resistant gene *Pm57* from *Aegilops searsii* into wheat. Theor Appl Genet. 2017; 130: 841–848.

39. Du P, Zhuang LF, Wang YZ, Yuan L, Wang Q, Wang DR, et al. Development of oligonucleotides and multiplex probes for quick and accurate identification of wheat and *Thinopyrum bessarabicum* chromosomes. Genome. 2016; 60: 93–103.

40. Li GQ, Fang TL, Zhang HT, Xie CJ, Li HJ, Yang T, et al. Molecular identification of a new powdery mildew resistance gene *Pm41* on chromosome 3BL derived from wild emmer (*Triticum turgidum* var. *dicoccoides*). Theor Appl Genet. 2009; 119: 531–539.

41. Li WT, Zhu ZW, Chern M, Yin JJ, Yang C, Ran L, et al. A natural allele of a transcription factor in rice confers broad-spectrum blast resistance. Cell. 2017; 170: 114–126.

42. Zhuang L, Sun L, Li AX, Chen TT, Qi ZJ. Identification and development of diagnostic markers for a powdery mildew resistance gene on chromosome 2R of Chinese rye cultivar Jingzhouheimai. Mol Breed. 2011; 27: 455–465.

43. Jiang J, Friebe B, Gill BS. Recent advances in alien gene transfer in wheat. Euphytica. 1993; 73: 199–212.

44. Zhang RQ, Fan YL, Kong LN, Wang ZJ, Wu JZ, Xing LP, et al. *Pm62*, an adult-plant powdery mildew resistance gene introgressed from *Dasypyrum villosum* chromosome arm 2VL into wheat. Theor Appl Genet. 2018; 131: 2613–2620.

45. Tan GX, Liu K, Kang JM, Xu KD, Zhang Y, Hu LZ, et al. Transcriptome analysis of the compatible interaction of tomato with *Verticillium dahliae* using RNA-sequencing. Front Plant Sci. 2015; 6: 428.

46. Xiao J, Jin XH, Jia XP, Wang HY, Cao AZ, Zhao WP, et al. Transcriptome-based discovery of pathways and genes related to resistance against *Fusarium* head blight in wheat landrace Wangshuibai. BMC Genomics. 2013; 14: 197.

47. Hao YB, Wang T, Wang K, Wang XJ, Fu YP, Huang LL, et al. Transcriptome analysis provides insights into the mechanisms underlying wheat plant resistance to stripe rust at the adult plant stage. PLoS One. 2016; 11: e0150717.

48. Wang DF, Wang XB, Mei Y, Dong HS. The wheat homolog of putative nucleotide-binding site-leucine-rich repeat resistance gene *TaRGA* contributes to resistance against powdery mildew. Funct Integr Genomics. 2016; 16: 115–126.

49. Sánchez-Martín J, Steuernagel B, Ghosh S, Herren G, Hurni S, Adamski N, et al. Rapid gene isolation in barley and wheat by mutant chromosome sequencing. Genome Biol. 2016; 17: 221.

50. Yahiaoui N, Srichumpa P, Dudler R, Keller B. Genome analysis at different ploidy levels allows cloning of the powdery mildew resistance gene *Pm3b* from hexaploid wheat. Plant J. 2004; 37: 528–538.

51. Hurni S, Brunner S, Buchmann G, Herren G, Jordan T, Krukowski P, et al. Rye *Pm8* and wheat *Pm3* are orthologous genes and show evolutionary conservation of resistance function against powdery mildew. Plant J. 2013; 76: 957–969.

52. Xing LP, Hu P, Liu JQ, Witek K, Zhou S, Xu JF, et al. *Pm21* from *Haynaldia villosa* encodes a CC-NBS-LRR protein conferring powdery mildew resistance in wheat. Mol Plant. 2018; 11: 874–878.

53. Du WL, Wang J, Lu M, Sun SG, Chen XH, Zhao JX, et al. Characterization of a wheat-*Psathyrostachys huashanica* Keng 4Ns disomic addition line for enhanced tiller numbers and stripe rust resistance. Planta. 2014; 239: 97–105.

54. King I, Purdie K, Rezanoor H, Koebner R, Miller T, Reader S, et al. Characterization of *Thinopyrum bessarabicum* chromosome segments in wheat using random amplified polymorphic DNAs (RAPDs) and genomic in situ hybridization. Theor Appl Genet. 1993; 86: 895–900.

55. Song LQ, Lu YQ, Zhang JP, Pan CL, Yang XM, Li XQ, et al. Physical mapping of *Agropyron cristatum* chromosome 6P using deletion lines in common wheat background. Theor Appl Genet. 2016; 129: 1023–1034.

56. Li SJ, Lin ZS, Liu C, Wang K, Du LP, Ye XG. Development and comparative genomic mapping of *Dasypyrum villosum* 6V#4S-specific PCR markers using transcriptome data. Theor Appl Genet. 2017; 130: 2057–2068.

57. Gupta P, Rustgi S. Molecular markers from the transcribed/expressed region of the genome in higher plants. Funct Integr Genomics. 2004; 4: 139–162.

58. Zhang HY, Wei LB, Miao HM, Zhang TD, Wang CY. Development and validation of genic-SSR markers in sesame by RNA-seq. BMC Genomics. 2012; 13: 316.

